# The asymmetric ABA response on both sides of root tip is important for tomato root hydrotropism by mediating proton efflux and cell elongation

**DOI:** 10.1101/2022.03.11.483958

**Authors:** Wei Yuan, Jianping Liu, Hui Dai, Qian Zhang, Weifeng Xu, Jianhua Zhang, Ying Li

**Author notes:** These authors contributed equally to this work (Wei Yuan, Jianping Liu). Corresponding author: Weifeng Xu,; Ying Li.

## Abstract

Hydrotropism is an important adaptation of plant roots to the uneven distribution of water, and the current research on hydrotropism is mainly focused on *Arabidopsis thaliana*. We examined hydrotropism in tomato (*Solanum lycopersicum*) primary roots. We used RNA sequencing to detect the gene expression on both sides (dry and wet side) of root tips (5 mm from the root cap junction) by splitting root tips longitudinally into two halves. We found that hydrostimulation induced the asymmetric cell elongation between the dry side (lower water potential) and wet side of root tips (higher water potential). ABA biosynthesis gene *ABA4* was induced on the dry side as compared to the wet side of root tips. Chemical inhibitors that block ABA biosynthesis can disrupt hydrotropism, and ABA biosynthesis mutant *not* showed significantly reduced hydrotropism. Furthermore, asymmetric H^+^ efflux was found in wild-type but not in root tips of ABA biosynthesis mutant *not* after hydrostimulation. Our results suggest that the asymmetric ABA response on both sides of root tip mediate asymmetric H^+^ efflux, and then drive the asymmetric cell elongation, which allows the root to bend towards the wet side to take up more water.

**Graphical abstract:** Involvement of ABA-mediated asymmetric H^+^ efflux in root hydrotropism.
Compared to the wet side of root tip (higher water potential), the dry side (lower water potential) induces the expression of ABA biosynthesis gene *ABA4*, thus enhancing proton efflux to promoting cell elongation on the dry side. Because H^+^ efflux and cell elongation on the dry side of the root tip are higher than that on the wet side, the asymmetric growth of cells on both sides allows the root to bend towards the wet side for taking up more water.

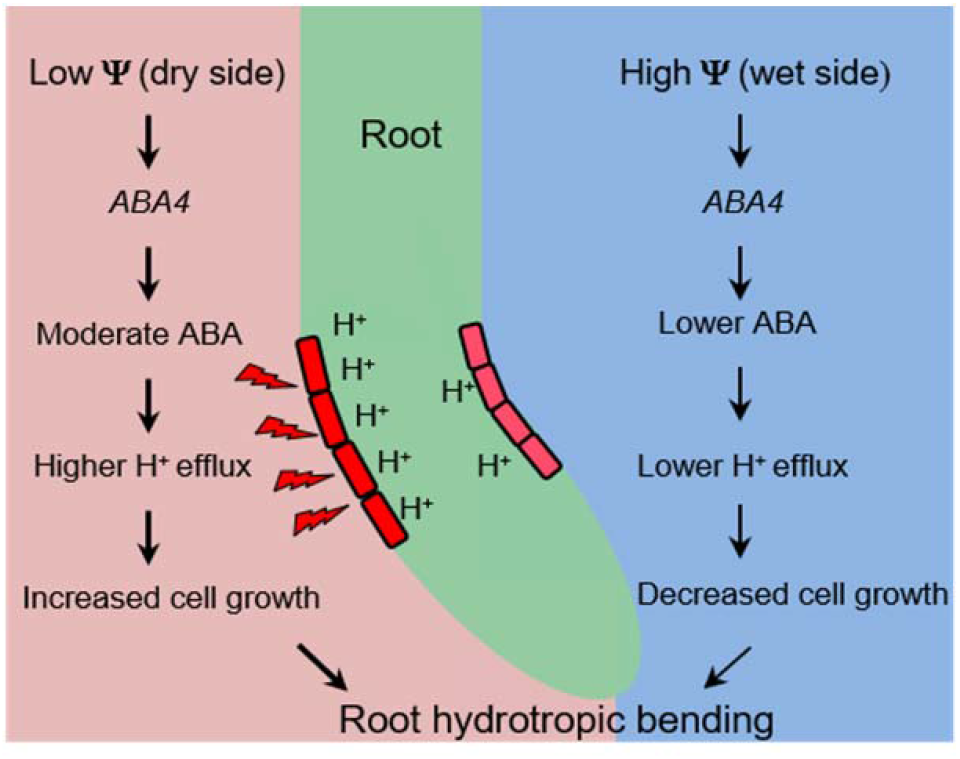

**In Brief:** The asymmetric ABA response on both sides of root tip (dry side and wet side) sequently mediates root asymmetric H^+^ efflux, and then drives the asymmetric cell elongation on both sides of root tip, which allows the root to bend towards the wet side for absorbing more water.

**Highlights:** Hydrotropic bending requires asymmetric cell elongation on the root two sides.

Asymmetric expression of ABA biosynthesis gene *ABA4* is required for root hydrotropism.

The H^+^ efflux on the dry side of the root is increased.

ABA-associated asymmetric H^+^ efflux driven root hydrotropic bending.

## Introduction

Water shortage is the major threat to agricultural production on a worldwide scale. Even in areas with sufficient water, the uneven distribution of water in its microenvironment requires continuous adaptation of plants to obtain more water (Chang et al., 2019; Li et al., 2020 a), that is called root hydrotropism: the plant roots sense the water potential gradient in its microenvironment and directed growth toward the moist soil (Takahashi et al., 2002; Li et al., 2020 b). Despite the significance of hydrotropism, our understanding of its physiological and molecular processes is very limited.

Abscisic acid (ABA) is a critical stress phytohormone, and the ABA signaling transduction pathway (PYR/PYL/RCAR-PP2Cs-SnRK2s) plays a pivotal role in coordinating the responses to decreased water availability (Sharp et al., 1994; Antoni et al., 2013; Rowe et al., 2016; Dietrich et al., 2017). Under stress conditions, ABA is rapidly induced and binds to Pyrabactin resistance1/PYR1-like/regulatory components of ABA receptor (PYR/PYL/RCAR) proteins, which subsequently repress group A PROTEIN PHOSPHATASES 2Cs (PP2Cs) (Ma et al., 2009; Nishimura et al., 2009). Concurrently, subclass III sucrose non-fermenting-1 related protein kinase 2 (SnRK2s) are released from PP2C-SnRK2 complexes to phosphorylate and activate a subgroup of the basic leucine zippers (bZIPs) transcription factors including ABA insensitive 5 (ABI5) and ABFs/ AREBs that recognize the ABRE promoter element (consensus PyACGTGG/TC) in ABA-responsive genes (Uno et al., 2000; Furihata et al., 2006; Fujii et al., 2007; Ma et al., 2009; Park et al., 2009). It has been suggested that ABA is required for root hydrotropism in *Arabidopsis* (Takahashi et al., 2002; Dietrich et al., 2017; Miao et al., 2020). The Arabidopsis *aba1-1* mutants were less sensitive to hydrostimulation, while application of ABA to *aba1-1* restored the normal sensitivity to the hydrotropic stimulation (Takahashi et al., 2002). The *no hydrotropic response 1* (*nhr1*) mutant had reduced root growth sensitivity to ABA (Eapen et al., 2003), and ABA up-regulates the expression of *MIZU-KUSSEI 1 (MIZ1*), a gene essential for root hydrotropism (Moriwaki et al., 2012). *112458*, a sextuple ABA receptor *PYR/PYL* mutant, displayed a reduced hydrotropism, whereas *Qabi2-2* plants, a *PP2Cs* quadruple mutant, exhibited enhanced hydrotropism in *Arabidopsis* (Antoni et al., 2013). Recent studies found that the SnRK2s protein kinases regulate hydrotropic response in cortical cells of the elongation zone (Dietrich et al., 2017). These findings suggest that hydrotropism depends on the core ABA signal transduction pathway.

Plasma membrane (PM) H^+^-ATPase (PM H^+^-ATPase), a subfamily of P-type H^+^-ATPases, generates a membrane potential and H^+^ gradient across the PM, energising various ion channels and multiple H^+^-coupled transporters for diverse physiological processes (Moloney et al., 1981; Hager, 2003; Falhof et al., 2016). Recently, we reported that brassinosteroid (BR) insensitive 1 (BRI1, a BR receptor) targets *Arabidopsis* plasma membrane (PM) H^+^-dependent adenosine triphosphatase (ATPase) 2 (AHA2) to mediate hydrotropic response in *Arabidopsis*, and the *bri1-5* mutant showed reduced root hydrotropism, which correlates with lower apoplastic H^+^ extrusion (Miao et al., 2018). More recently, we found that ABA-insensitive 1 (ABI1), a key component of PP2C in ABA signal pathway, interacts directly with the C-terminal R domain of AHA2 and dephosphorylates its penultimate threonine residue (Thr^947^), which decreases PM H^+^ extrusion and negatively regulates root hydrotropic response in *Arabidopsis* (Miao et al., 2020).

In this study, we found that ABA-mediated asymmetric H^+^ efflux driven root hydrotropic bending in tomato. The H^+^ efflux, the expression of ABA biosynthesis gene *ABA4* and cell elongation on the dry side of the tomato root tips (5 mm from the root cap junction) are higher than that on the wet side, the asymmetric growth of cells on both sides of the root allows the root to bend towards the wet side to take up more water. These findings disclose a novel opportunity for breeding tomato varieties with improved water-use efficiency under water-limited conditions.

## Materials and Methods

### Plant materials and growth conditions

Four tomato (*Solanum lycopersicum*) cultivars including Micro-Tom (MT), LA0534 (LU), OFSN (OFSNOFSN), and Ailsa Craig (AC, LA2838) were used for the analysis of hydrotropism. The background of ABA-deficient mutant *notabilis (not*, LA0617) is LU. Tomato seeds were surface sterilized with 30% sodium hypochlorite (NaClO) and distilled water at the volume ratio of 1:2 for 3 minutes, and then rinsed with sterile distilled water for 5 times. Add appropriate amount of sterile distilled water to the sterilized tomato seeds, and then soak them at 30°*C* in the dark for 2 days to make sure the seeds fully absorb water. Subsequently, the seeds were sown on 1% agar containing 1/2 Murashige and Skoog (1/2 MS) media at 22°*C* under 16 h light/ 8 h dark photoperiod. Five-day-old uniform seedlings were used for subsequent experiments.

### Root hydrotropism assays

The agar-sorbitol system shown in Figure S1 was established as previously described (Takahashi et al., 2002). First, about 50 ml of 1/2 MS medium was pour into a 13 ×13 cm square dish. After coagulation, cut off the lower left of the medium with a blade (with an inclination of about 57 degrees, 2 cm from the upper and lower boundaries), and pour about 25 ml of 1/2 MS with sorbitol medium, the two different medium are on the same level as far as possible. A water potential gradient was formed from lower left (low water potential) to upper right (high water potential) (Figure S1). The uniform tomato seedlings were vertically moved to the normal 1/2 MS medium, and the root tip was 3 mm above the boundary of the two media. Each seedling was separated by about 1.0 cm, sealed with sealing film, and placed in a growth chamber. The hydrotropic bending and elongation of roots were photographed using a digital camera (Nikon D7100) and were measured using ImageJ software.

### Treatment with exogenous ABA and ABA inhibitor Fluridone

Exogenous ABA and ABA inhibitor Fluridone (FLU) were prepared at 1000 × concentration. Exogenous ABA or FLU was added into 50 ml 1 / 2 MS medium. After coagulation, cut half of the medium (the method is the same as above), and pour into about 25 ml 1/2 MS + 1000 mm sorbitol medium with a certain amount of ABA or FLU.

### Bromocresol purple dyeing

The pH-sensitive indicator bromocresol purple (pH range is 5.2 (yellow) to 6.8 (purple))was used as described (Bissoli et al., 2012). Briefly, tomato plants grown on control and hydrostimulation condition for 3 h were transferred to 1/2 MS vertical plates with 0.003% bromocresol purple and incubated in the light for 6-8 h. For this measurement, density of the stained areas was measured using ImageJ software. The density of control was taken to be 100%, and relative density was calculated based on control levels.

### Confocal microscopy

The root tips of tomato seedlings under control conditions or hydrostimulation for 3 h were stained with propidium iodide (PI, 40 μg/mL) dye solution. After 3 h of hydrostimulation, the slightly curved root tips were placed in PI dye solution on the slide with tweezers to keep the direction of the left and right sides of the root consistent with the growth direction on the hydrostimulation medium, and the cover glass was covered. The fluorescence of PI in tomato root tips was observed with a Zeiss LSM 780 laser spectral scanning confocal microscope as described by Xu *et al*. (Xu et al., 2013). PI was excited at 553 nm and detected at 615 nm.

The root tips of tomato seedlings under normal conditions or hydrostimulation for 3 h were stained with 8-hydroxypyrene-1,3,6-trisulfonic acid trisodium salt (HPTS) (Han and Burgess, 2010; Barbez et al., 2017). The samples were soaked in 50 mm HPTS dye solution about 30 minutes, and then the samples were washed with distilled water for 3 times. the method is the same as PI. For HPTS, excitation at 405 nm (protonation) and 458 nm (deprotonation) and emission at 514 nm. Divide the signal intensity of 458 nm channel of each pixel by the signal intensity of 405 nm channel to get a ratio image. The intensity value is related to the proton secretion of cell wall. All images were taken under identical conditions. For this measurement, signal density was measured using ImageJ software. The density of control were taken to be 100%, and relative density was calculated based on control levels.

### RNA sequencing and data analysis

The root tips of tomato (5 mm from the root cap junction) were obtained after 3 h of control or hydrostimulation; and the dry and wet sides of the tomato root tip (5 mm from the root cap junction) after 3 h of hydrostimulation were obtained by splitting longitudinally whole root into two halves. Three biological replicate samples (a total of 12 samples) were collected: the 5 mm of root tips grown in control conditions: Control 1, Control 2, Control 3; the 5 mm of root tips grown in hydroptimulated conditions: Hydroptimulated 1, Hydroptimulated 2, Hydroptimulated 3; the dry sides of the 5 mm of root tips grown in hydroptimulated conditions: Dry1, Dry 2, Dry 3; the wet sides of the 5 mm of the root tips grown in hydroptimulated conditions: Wet 1, Wet 2, Wet 3 (Figure 2). Each sample was collected into a 1.5 ml centrifuge tube without RNase, and quickly put into liquid nitrogen, and then transferred to −80 *°C* for storage.

**Figure 1.**
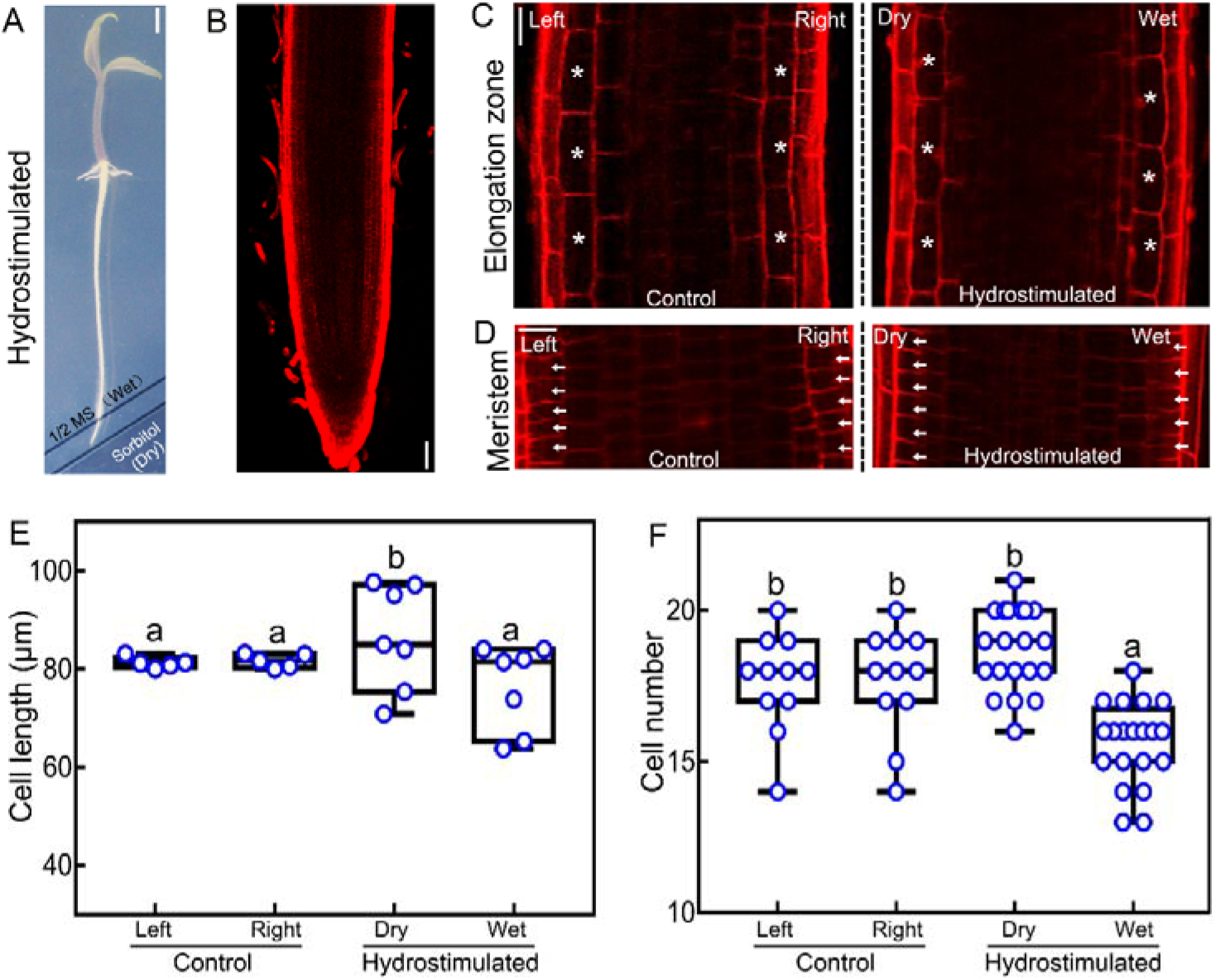
Differential cell elongation and cell number of both sides (dry side and wet side) in tomato root tip under hydrostimulation. (A) Representative images of tomato roots hydrostimulated for 8 h. Scale bar: 3 mm. (B) Representative micrographs of tomato roots hydrostimulated for 3 h. Roots were stained with PI. Scale bar: 100 μm. (C and D) The cell length on both sides in elongation zone (C) or cell number in meristematic zone (D) of tomato root tip under control or hydrostimulated for 3 h. Control roots grew on normal 1/2 MS conditions without water potential gradient, while hydrotimulated roots grew on agar-sorbitol system with water potential gradient. Dry, the side facing sorbitol; Wet, the side facing normal 1/2 MS. Roots were stained with PI. Asterisks (C) denote cell on both sides of elongation zone, and arrows (D) denote signal cell on both sides of meristematic zone. Scale bar: 40 μm. (E and F) Quantification of cell length (E) in elongation zone or cell number in meristematic zone (F) in the roots of plants described in (C) and (D). Data in E and F are presented as means ± SE of three independent biological replicates; different letters denote significant differences (*P*<0.05, Duncan’s test).

**Figure 2.**
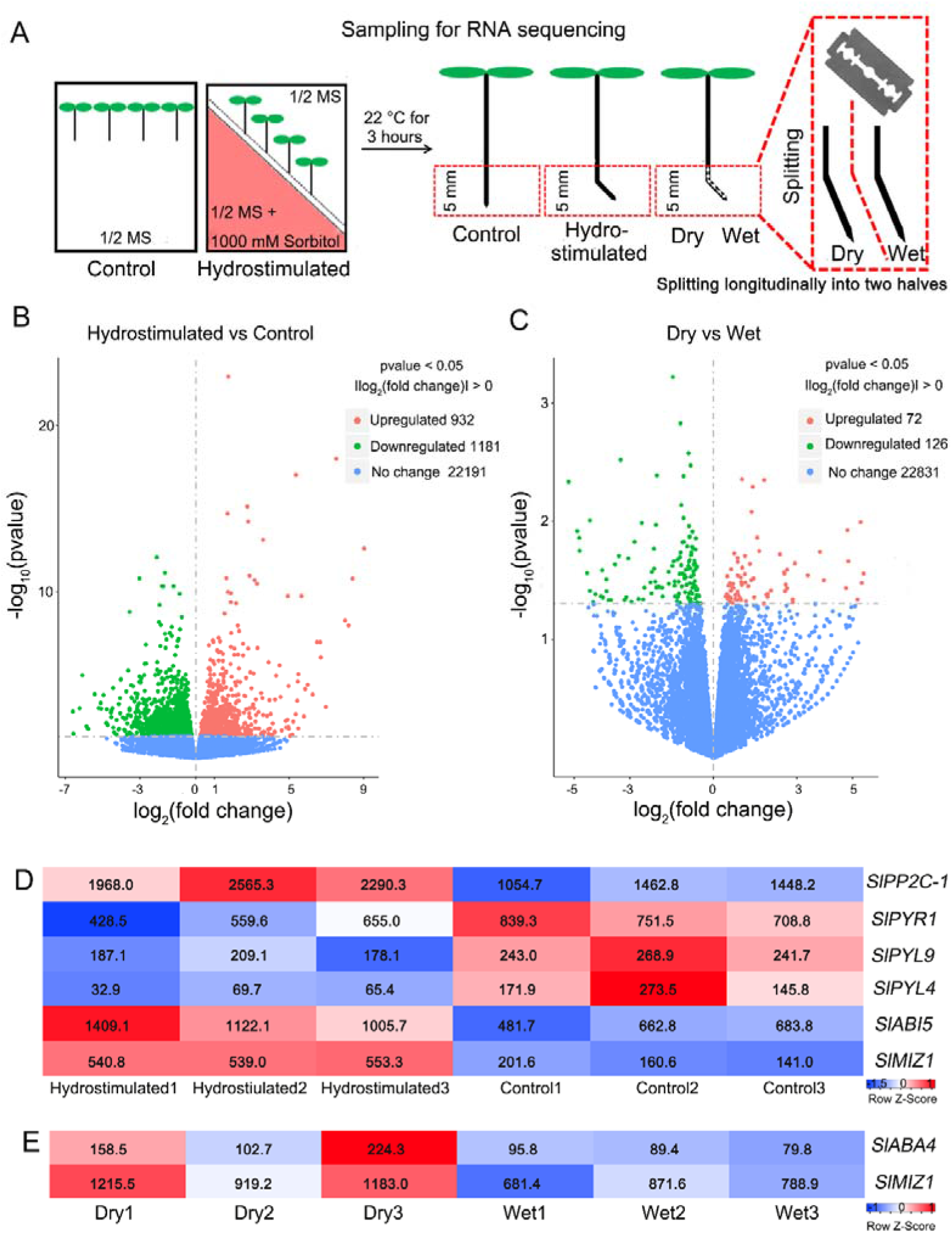
Response of ABA pathway in 5 mm area of both sides (dry side and wet side) in tomato root tips under hydrostimulation. ABA-related genes are differentialy expressed in 5 mm area (5 mm from the root cap junction) of tomato root tips under hydrostimulation. (A) Flow chart of RNA sequencing for harvesting samples in 5 mm area and dry / wet side of 5 mm area of tomato root tips. Control roots are the root tips 5 mm area (5 mm from the root cap junction) under normal 1/2 MS conditions; hydrotimulated roots are the root tips 5 mm area (5 mm from the root cap junction) under hydrostimulation; dry is the side of 5 mm root tips facing sorbitol under under hydrostimulation; and wet is the side of 5 mm root tips facing normal 1/2 MS under hydrostimulation. (B and C) Relationship between average expression of control vs hydrostimulated (B) and dry vs wet (C) and fold change for each gene. Each dot in the graphs represents a single gene, and red represents upregulated differentially expressed genes (DEGs), green represents downregulated DEGs, and blue represents no change. (D and E) Heatmap visualizing the expression patterns of DEGs in the ABA-related pathway under hydrostimulation. Each row represents one gene, columns represent the different treatments, and low expression levels are in blue and high expression levels are in red. PP2C, protein phosphatase 2C; PYR, pyrabactin resistance1; PYL, PYR1-like; ABI5, ABA insensitive 5; ABA4, ABA deficient 4; MIZ1, MIZU-KUSSEI 1.

The total RNA extraction, library construction and sequencing of samples were was performed by Novogene Co., LTD (Beijing Nuohe Zhiyuan Technology Co., LTD). RNA integrity was assessed using the RNA Nano 6000 Assay Kit of the Bioanalyzer 2100 system (Agilent Technologies, CA, USA). Sequencing libraries were generated using NEBNext UltraTM RNA Library Prep Kit for Illumina (NEB, USA) following manufacturer’s recommendations and index codes were added to attribute sequences to each sample. Raw data (raw reads) of fastq format were firstly processed through in-house perl scripts. In this step, clean data (clean reads) were obtained by removing reads containing adapter, reads containing ploy-N and low quality reads from raw data. The RPKM method (reads per kilobase of transcript per million mapped reads) was used to determine transcript abundance. Differential expression analysis of two conditions/groups (three biological replicates per condition) was performed using the DESeq2 R package (1.16.1). Genes with an adjusted P-value <0.05 found by DESeq2 were assigned as differentially expressed.

### qRT-PCR analysis

Total RNA was extracted from tomato roots using a plant RNA Kit (OMEGA, USA) according to the manufacturer’s instructions. First-strand cDNA was synthesized using a *TransScript* One-Step gDNA Removal and cDNA Synthesis SuperMix Kit (TransGen Biotech, China) according to the manufacturer’s instructions. Analyses with qRT-PCR were performed using a *TransScript* Tip Green qPCR SuperMix Kit (TransGen Biotech, China) and a CFX96 Real-time PCR Detection system (Bio-Rad, USA) according to the manufacturers’ instructions. The specific primers for each gene are listed in Table S1. Results were normalized using *α-Tubulin* gene as the endogenous control.

### Statistical analysis

For all experiments, statistical tests were carried out using SPSS software (IBM Corporation, USA). A two-tailed Student’s *t*-test was used to compare between two groups. For comparisons between more than two groups, Duncan’s test was used. Data were represented as the mean ± SEM from at least three independent experiments, and differences were considered significant at p < 0.05.

## Results

### Hydrotropic bending involves differential cell elongation on dry and wet sides of tomato roots

The 400 mM sorbitol induced-water potential gradients (agar-sorbitol system) is commonly employed to observe the root hydrotropism in model plant *Arabidopsis thaliana* (Takahashi et al., 2002). To find the most suitable experimental system for the study of tomato root hydrotropism, we used different water potential gradients formed by different concentations of sorbitol (400 mM, 800 mM, 1000 mM, 1200 mM and 1500 mM) to observe the root hydrotropic bending and root growth of tomato (Micro-Tom) (Figure S1 and S2). Results showed that hydrotropism of Micro-Tom roots in higher concentration of sorbitol (800 mM, 1000 mM, 1200 mM and 1500 mM) was higher than that in the 400 mM sorbitol treatment (Figure S2A). In addition, we found that the root growth was severely inhibited when the tomato Micro-Tom was treated under the water potential gradient formed by the 1200 mM and 1500 mM sorbitol treatments (Figure S2B). However, the root elongation was not seriously inhibited under the water potential gradient formed by 1000 mM sorbitol treatment, and the hydrotropic bending of tomato roots was stronger (Figure S2). Therefore, the water potential gradient formed by 1000 mM sorbitol was the most suitable system to study the hydrotropism of tomato roots. Next, we compared the hydrotropic responses of different tomato ecotypes (Micro-Tom、 LU、 OFSN、 AC) using the 1000 mm sorbitol-induced water potential gradient. Results showed that LU exhibited significantly higher hydrotropism (Figure S3). Therefore, we used LU for subsequent experiments.

To explore the mechanism of tomato root hydrotropic bending (Figure 1A), we observed the cell growth of meristem zone and elongation zone by PI staining (Figure 1B-D). Under control conditions, there was no significant difference in cell length between the left and right sides of the elongation zone, and the cell length on both sides showed symmetrical growth (Figure 1C). However, the cells in the elongation zone exhibited asymmetric growth between the dry and wet sides under hydrostimulation, and the cell length on the dry side was more longer than that on the wet side (Figures 1C and 1E). Moreover, we also found that there was no significant difference on the cell numbers between the left and right sides of the root apical meristem zone under control conditions, while the cell numbers on dry side were significantly more than that of wet side under hydrostimulation (Figures 1D and 1F). Taken together, these results suggest that the asymmetrical growth of tomato root tips between dry side and wet sides drive hydrotropic bending.

### Hydrostimulation induces asymmetric expression of ABA-related genes in tomato roots

Hydrotropic bending involves asymmetrical growth of cells on both sides of root tips, and we first thought of using single-cell RNA-sequencing (scRNA-seq) to detect the gene expression changes of cells on both sides of root tips by splitting root tips longitudinally into two halves. However, high quality cells on both sides of root tips were not obtained successfully. Thus, we employed transcriptomic analyses (RNA-sequencing) to identify the genes that are involved in early hydrotropic response of dry side- and wet side-exposed roots (Figure 2, Figure S4). The samples included: the 5 mm of root tips (5 mm from the root cap junction) grown under control conditions (Control); the 5 mm of root tips grown under hydrostimulation conditions (Hydrostimulated); the 5 mm dry side of root tips grown under hydrostimulation conditions (Dry); and the 5 mm wet side of root tips grown under hydrostimulation conditions (Wet) (Figure 2A, Figure S4). RNA sequencing analysis was conducted using three biological replicates of each treatment. In total, 12 libraries were constructed and analyzed (Figure 2A, Figure S4). Correlation heatmap analysis showed that pearson correlation among three biological replicates was higher (Figure S4A). Furthermore, we found that hydrostimulation-induced differentially expressed genes (DEGs) were significantly different between dry side and wet side in tomato roots (Figure S4B).

There were 2,113 DEGs between control and hydrostimulation and 198 DEGs between dry side and wet side under hydrostimulated treatment (Figures 2B and 2C, Figure S5). Furthermore, compared with the wet side, 126 genes were significantly upregulated in the dry side, while 72 genes were significantly downregulated (Figure 2B and 2C, Figure S5). Among these, a number of genes that are associated with phytohormone abscisic acid (ABA) (Figures 2D and 2E, Figure 3A). In plants, the first step of the ABA biosynthesis pathway involves a conversion of zeaxanthin using zeaxanthin epoxidase (ZEP/ABA1) as the enzyme (Figure 3A) (Nambara and Marion-Poll, 2005). Subsequently, *all*-*trans*-Violaxanthin is converted into *all-trans* Neoxanthin by ABA DEFICIENT4 (ABA4), and then 9-cis-epoxycarotenoid dioxygenases (NCEDs) converts 9-cis-Neoxanthin into xanthoxin, which is further converted into ABA by ABA DEFICIENT2 (ABA2), ABA aldehyde oxidase 3 (AAO3) and ABA DEFICIENT3 (ABA3) (Nambara and Marion-Poll, 2005). In the presence of ABA, ABA binds to the ABA receptor PYR/PYL/RCAR proteins to impede the activity of protein phosphatases 2C (PP2Cs) that inhibit the activity of SnRK2 protein kinases, and then the SnRK2s are released from PP2C-dependent negative regulation, allowing SnRK2s to regulate downstream transcription factors (TFs), including bZIP transcription factors ABA INSENSITIVE 5 (ABI5) and ABA-responsive element (ABRE)-binding protein/ABRE binding factors (AREB/ABFs), activating the expression of many ABA-responsive genes in response to dehydration stress in plants (Uno et al., 2000; Furihata et al., 2006; Fujii et al., 2007; Ma et al., 2009; Park et al., 2009). In this study, genes related to ABA biosynthesis, signaling pathway and ABA-responsive genes were detected, and the expression of most of these genes was increased under hysrostimulation conditions (Figures 2D and 2E). In addition, a previously known hydrotropic gene (*MIZ1*) was induced by hydrostimulation, and the expression of which in the dry side was higher than that in the wet side (Figures 2D and 2E).

**Figure 3.**
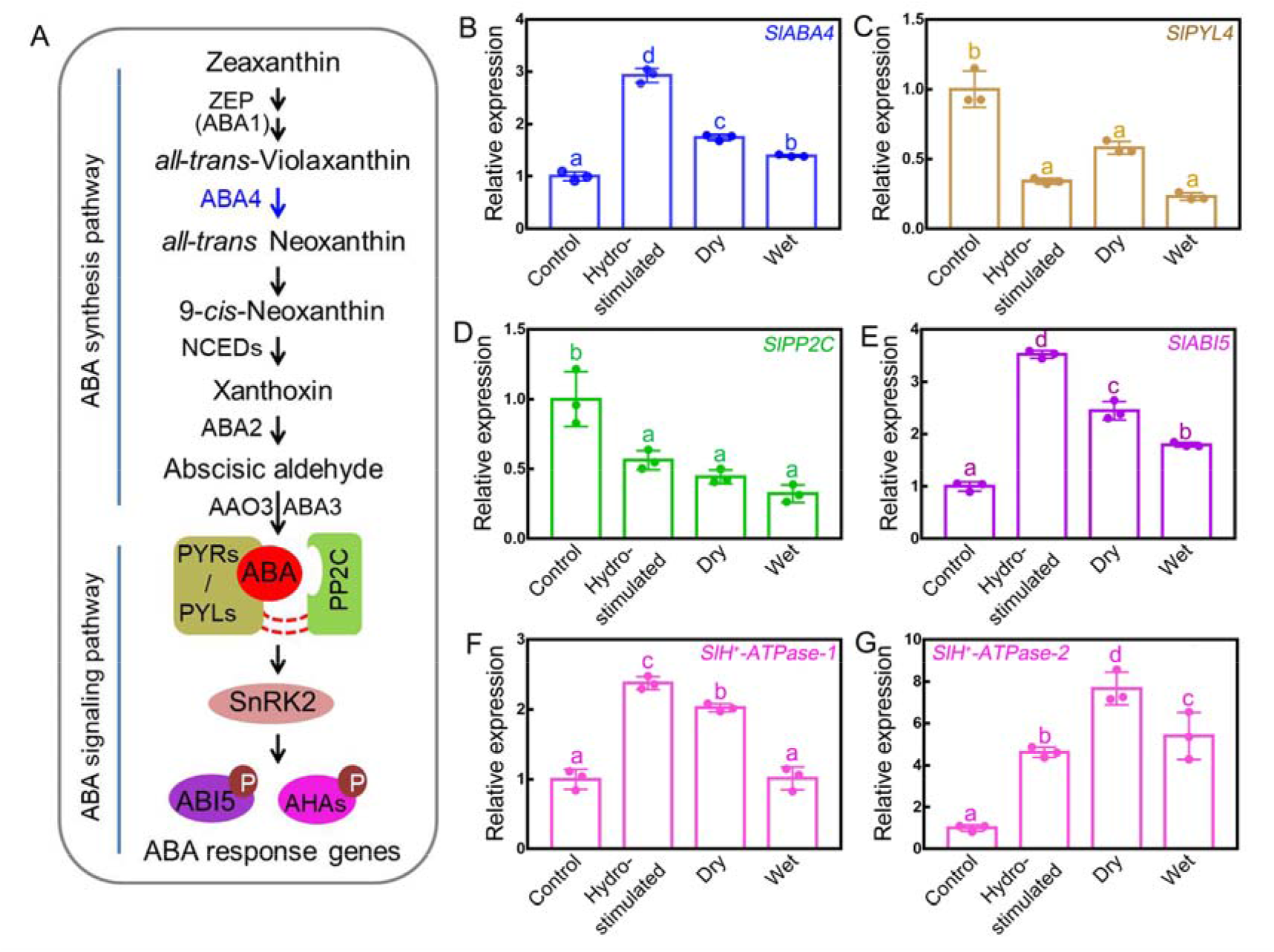
ABA-related genes are differentialy expressed in 5 mm area of both sides (dry side and wet side) in tomato root tips under hydrostimulation. (A) Schematic overview of ABA synthesis and signaling pathway. (B-G) The *SlABA4* expression (B), *SlPYL4* expression (C), *SlPP2C* expression (D), *SlABI5* expression (E), *SlH^+^-ATPase-1* expression (F) and *SlH^+^-ATPase-2* expression (G) in 5 mm area of tomato root tips under hydrostimulation. Genes expression (B-G) under control were taken to be 100%, relative genes expression under hydrostimulation were calculated on the basis of control levels. *a-Tubulin* was used as the internal control. Control roots are the root tips 5 mm area (5 mm from the root cap junction) under normal 1/2 MS conditions; hydrotimulated roots are the the root tips 5 mm area (5 mm from the root cap junction) under hydrostimulation; dry is the side of 5 mm root tips facing dry sorbitol under under hydrostimulation; and wet is the side of 5 mm root tips facing normal 1/2 MS under hydrostimulation. Data in B-G are presented as means ± SE of three independent biological replicates; different letters denote significant differences (*P*<0.05, Duncan’s test).

Quantitative Real time-PCR (qRT-PCR) was used to further detect the expression of ABA-related genes in the root tip 5 mm area (5 mm from the root cap junction) or dry side of 5 mm root tips facing dry sorbitol and wet side of 5 mm root tips facing normal 1/2 MS under hydrostimulation (Figures 3B-G). The results showed that the expression of *ABA4* (ABA synthesis-related gene) and *ABI5* (downstream transcription factor of ABA signaling pathway) were significantly increased under hydrostimulated treatment than that in control conditions, and the expression levels of *ABA4* and *ABI5* in dry side were also significantly higher than those on the wet side (Figures 3B and 3E). The expression pattern of *MIZ1* was similar to *ABA4* (Figure S6). The expression levels of *PYL4* (a member of ABA receptor family) and *PP2C* (a negative regulator of the ABA signaling) were lower than normal conditions (Figures 3C and 3D). In addition, the expression of plasma membrane H^+^-ATPase (proton pump) genes after hydrostimulated treatment was significantly increased compared with control, and the expression of H^+^-ATPase genes in dry side was significantly higher than that in wet side (Figures 3F and 3G). The above results suggested that asymmetric expression of ABA-related genes may be an important reason for the root hydrotropism in tomato.

### ABA-mediated H^+^ efflux plays an important role in hydrotropism of tomato roots

To further verify the role of ABA in tomato root hydrotropism, we added 1 μM ABA and 10 μM Fluridone (FLU, an ABA inhibitor) to observe the root hydrotropism. The results showed that the hydrotropism was slightly increased after 1 μM ABA treatment, while 10 μM FLU significantly inhibited the root hydrotropism (Figure 4A). Then, we examined the hydrotropism of tomato ABA-deficient mutant *not*. As shown in Figure 4B, the hydrotropic response of wild-type LU was nearly 30 degrees, while the hydrotropic response of *not* mutant was about 15 degrees. These results further confirmed that ABA positively regulates root hydrotropism in tomato.

**Figure 4.**
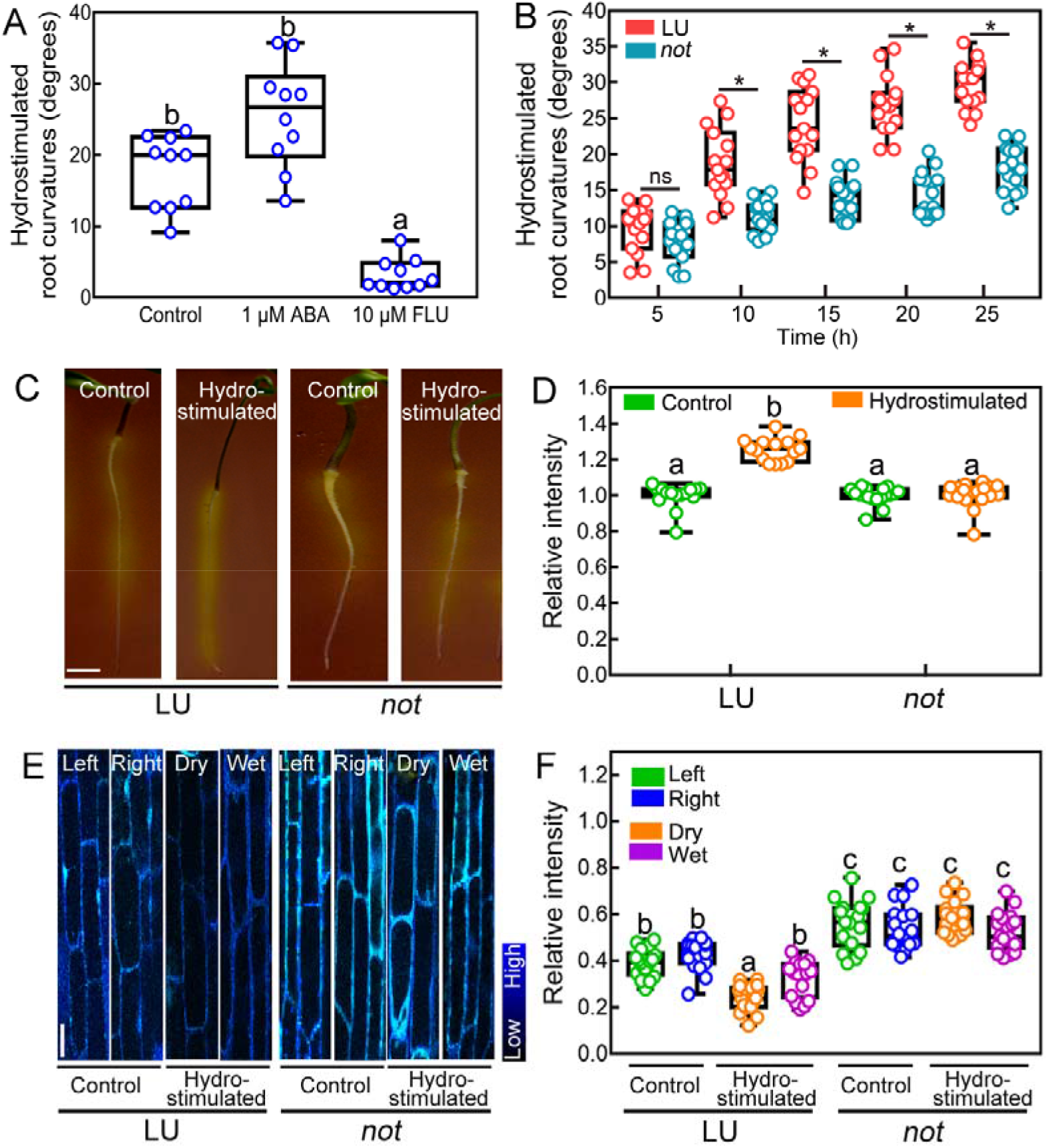
ABA-mediated asymmetric H^+^ efflux positively regulates the hydrotropism of tomato root tips. (A) Hydrotropic bending of tomato wild-type (LU) roots under different treatments: 1 μM ABA and 10 μM FLU (fluridone, ABA biosynthetic inhibitor). (B) Hydrotropic bending of tomato LU and ABA deficient mutants *not*. (C) Representative images of LU and ABA deficient mutant *not* roots stained with pH indicator bromocresol purple after 3 h control or hydrostimulation. A yellow colour around the roots indicates proton (H^+^) extrusion. Scale bar: 6 mm. (D) Quantification of proton (H^+^) extrusion in the root of plants described in (C). (E) Representative images for HPTS staining of tomato LU and ABA deficient mutants *not* apoplastic epidermal cells in root elongation zone after 3 h control or hydrostimulation. Control roots grew on normal 1/2 MS conditions without water potential gradient, while hydrotimulated roots grew on agar-sorbitol system with water potential gradient. Dry, the side facing dry sorbitol; Wet, the side facing normal 1/2 MS. Scale bar: 25 μm. Color code (black to blue) describes (low to high) 458/405 intensity and pH values. (F) Quantification of H^+^ efflux in the root of plants described in (E).The relative intensity correlate with the 458/405 ratio values. The higher the 458/408 ratio, the higher the pH and less apoplastic H^+^. Data in A, B, D and F are presented as means ± SE of three independent biological replicates; different letters denote significant differences (*P*<0.05, Duncan’ s test).

Previous studies and our recent study have suggested that ABA is closely related to H^+^ efflux (Hayashi et al., 2014; Miao et al., 2020; Planes et al., 2015). We next determined whether the H^+^ efflux is changed in root tip cells of tomato wild-type LU and ABA-deficient mutant *not* under control or hydrostimulated conditions by using the pH-sensitive dye bromocresol purple (acid-base indicator) (Figures 4C and 4D) and HPTS staining (a fluorescent pH-indicator) (Figures 4E and 4F). As shown in Figure 4C, the roots of wild-type LU were dyed deeper and secreted the most H^+^ under hydrostimulated conditions compared with controls, while the roots of *not* mutant under both control and hydrostimulated conditions were dyed shallower and secreted less H^+^. Then, we used Image J to quantify the protons secretion at whole root of LU and *not* mutant under control and hydrostimulated conditions (Figure 4D) or 5 mm root tip of LU and *not* mutant under control and hydrostimulated conditions (Figure S7). As shown in Figure 4D, the H^+^ secreted from the whole roots of wild-type LU was enhanced under hydrostimulated treatment, while the H^+^ secreted from the roots of ABA-deficient mutant *not* under hydrostimulated treatment was similar to control conditions. At the same time, the H^+^ secretion at 5 mm root tips of wild-type LU and mutant *not* under hydrostimulation was stronger than the respective controls, but the H^+^ secretion in wild-type LU in this region was stronger than that in *not* under hydrostimulation (Figure S7). Next, we introduced 8-hydroxypyrene-1,3,6-trisulfonic acid trisodium salt (HPTS) as a suitable fluorescent pH-indicator for assessing apoplastic pH in elongation zone cells in tomato roots (Barbez et al., 2017; Han and Burgess, 2010). The apoplastic pH correlates with the ratiometric values (the 458 nm signal intensity has been divided by the 405 nm intensity). The higher the 458/408 ratio means the higher the pH and less apoplastic H^+^ (Barbez et al., 2017). Under control conditions, there was no significant difference in elongation zone cells apoplastic H^+^ between the left and right sides of wild-type LU or *not* roots (Figures 4E and 4F). Under hydrostimulation, apoplastic H^+^ in the dry side (lower water potential) of wild-type LU roots was higher than the wet side, while apoplastic H^+^ in the dry side of *not* mutant was similar to the wet side (Figures 4E and 4F). These results indicated that ABA positively regulates asymmetric H^+^ efflux in the root tip, which enhances cell elongation to drive root hydrotropic bending.

## Discussion

At present, the hydrotropism research is mainly focused on *Arabidopsis thaliana* (Dietrich et al., 2017; Chang et al., 2019; Li et al., 2020a), and there are few studies on crop plants. In this study, we examined root hydrotropism in tomato plants. We found that the water potential gradients formed by 1000 mM sorbitol was the most suitable hydrotropism experimental system for the study of tomato plants (Figures S1 and S2). The hydrotropism of *Arabidopsis* roots is strong on the water potential gradient formed by 400 mM sorbitol (Takahashi et al., 2002). Because the root of tomato is thicker than that of *Arabidopsis*, it is understandable that tomato roots need a higher water potential gradient to bend towards more water.

Under hydrostimulation, the dry side (lower water potential) of the primary roots contain more meristematic cortex cells compared to the wet side (higher water potential) of the primary roots in tomato (Figures 1D and 1F). These results are consistent with previous studies on *Arabidopsis* showing that the increase of cell number in the meristem zone on the dry side leads to the root bending towards the wet side (Chang et al., 2019). Furthermore, the cells length of the dry side in the elongation zone was longer than that in the wet side, indicating that the cells on both sides of the root tip grew asymmetrically during hydrostimulation (Figures 1C and 1E). These results are consistent with previous studies showing that hydrotropic bending involves coordinated adjustment of spatial cell elongation and cell flux in Arabidopsis (*Arabidopsis thaliana*) (Dietrich et al., 2017; Chang et al., 2019) and maize (*Zea mays*) (Wang et al., 2020). Therefore, compared with the dry side, the wet side of the roots has more active cell division and faster cell growth, which leads to the root bending towards water.

Previous studies indicated that *Mizu-kussei 1* (*MIZ1*) is crucial for root hydrotropism, and MIZ1 contains an unknown domain (DUF617) with unknown function, which is called MIZ domain (Kobayashi et al., 2007). MIZ1 domain is conserved in terrestrial plants including bryophytes, and genes containing MIZ1 domain are called MIZ like genes and have similar functions (Kobayashi et al., 2007). Shkolnik recently reported that MIZ1 directly interacted with the endoplasmic reticulum (ER) Ca^2+^-ATPase isoform ECA1, causing an asymmetrical distribution of Ca^2+^ in the elongation zone of *Arabidopsis* roots prior to hydrotropic bending (Shkolnik et al., 2018; Chang et al., 2019) recently reported that MIZ1 is required for the formation of cytokinin asymmetric distribution and the expression levels of ARRs under hydrostimulation. Although it has been shown that the MIZ1 is crucial for responding to moisture gradients, there is no direct evidence for the asymmetric expression of *MIZ1* in roots under hydrostimulation (Moriwaki et al., 2010; Fujii et al., 2018). Our results revealed that the expression of *MIZ1* in tomato root tips was significantly up-regulated under hydrostimulation (Figures 2B and 2D; Figure S6), and more important, asymmetric expression of *MIZ1* was indeed detected in both sides of the 5 mm of tomato root tip under hydrostimulation by splitting root tips longitudinally into two halves (Figures 2C and 2E; Figure S6). These results indicated that asymmetric expression of *MIZ1* drives tomato root hydrotropic bending and provided further evidence for the important role of *MIZ1* in root hydrotropism.

It has been shown that the osmotic stress hormone ABA is required for root hydrotropism (Takahashi et al., 2002; Dietrich et al., 2017; Miao et al., 2020). However, there is no evidence for the asymmetric ABA responses across the root in response to moisture gradients. Our results showed that the core components of ABA signaling were differently expressed under hydrostimulation (Figure 2D), and more important, the ABA biosynthesis gene *ABA4* was indeed asymmetrically expressed in both sides of the 5 mm of tomato root tips under hydrostimulation (Figure 2E; Figure 3B), that is, the expression of *ABA4* in the dry side of the root were higher than that in the wet-side. In roots, whilst high ABA levels inhibit growth (Fujii et al., 2007), moderate ABA levels promote elongation by regulating PM H^+^-ATPase-mediated H^+^ efflux at low water potential (Janicka-Russak and Kłobus, 2007; Xu et al., 2013). In addition, our recent studies found that PM H^+^-ATPase-mediated H^+^ efflux is also crucial for responding to moisture gradients in *Arabidopsis thaliana* (Miao et al., 2018; Miao et al., 2020). In tomato plants, *PM H^+^-ATPases* were asymmetrically expressed in both sides of the 5 mm of tomato root tips under hydrostimulation (Figures 3F and 3G), and more H^+^ efflux was observed after hydrostimulation treatment, and ABA deficient mutant *not* showed lower H^+^ efflux than wild-type LU (Figures 4C and 4D; Figure S7). Furthermore, asymmetric apoplastic H^+^ was found in wild-type LU but not in ABA biosynthesis mutant *not* after hydrostimulation treatment (Figures 4E and 4F). H^+^ can acidify the cell wall in plants, and cell wall acidification triggers cellular elongation (Moloney et al., 1981; Hager, 2003; Falhof et al., 2016). Thus, the asymmetric apoplastic H^+^ was positively correlated with asymmetric cell growth (Figures 4E and 4F; Figures 1C-F). Taken together, these results suggest that differential ABA response positively regulates asymmetric H^+^ efflux in the root tip, which enhances cell elongation and drive root hydrotropic bending.

In conclusion, our results suggest that ABA-mediated asymmetric H^+^ efflux is crucial for root hydrotropism. The expression of ABA-related genes and H^+^ efflux on the dry side of the root are increased under hydrostimulation, and the enhanced H^+^ efflux promotes cell elongation and drive root hydrotropic bending. Uncovering the detailed physiological and molecular mechanisms of hydrotropism can help us to carry out novel strategies to generate tolerant crop plants with higher water-use efficiency and higher productivity under drought environmental conditions.

## Acknowledgements

We are grateful for grant support from the National Natural Science Foundation of China (31872169, 31761130073 and 31422047).

## Author Contribution

Y. L., W.Y. and W.X. planned and designed the research, Y. L. and H.D. performed experiments and analysed data. Y. L., W.Y., J.P., J. Z. and W.X. wrote the manuscript. W.X. agrees to serve as the author responsible for contact and ensures communication.

## Supporting Information

